# Virtual Brain Modelling Reveals Altered Network Dynamics and Predicts Symptom Severity in Schizophrenia: Relevance to Antipsychotic Treatment Response and Resistance

**DOI:** 10.64898/2026.10.13.758782

**Authors:** Matteo Vitacca, Doris Pischedda, Mariateresa Ciccarelli, Giuseppe Pontillo, Giuseppe De Simone, Sirio Cocozza, Filip Ćipranić, Anita Monteverdi, Paolo Fusar-Poli, Claudia Angela Michela Gandini Wheeler-Kingshott, Fulvia Palesi, Andrea de Bartolomeis, Egidio D’Angelo

**Author notes:** Corresponding authors: Matteo Vitacca, Neuroimaging and Brain Modelling laboratory, Via Ferrata 5, 27100, Pavia, Egidio D’Angelo, Neurophysiology Unit, Via Forlanini 6, 27100 Pavia. These authors contributed equally to this work (Matteo Vitacca, Doris Pischedda).

## Abstract

**Background:** Schizophrenia (SZ) is a heterogeneous disorder characterized by positive, negative, cognitive and disorganisation symptoms, with pleomorphic phenotype and poorly understood aetiology. Dopamine dysregulation represents the most replicated in vivo pathophysiology finding. However, accumulating evidence suggests altered large-scale brain connectivity and excitation/inhibition balance disruption. While quantifying these alterations would help clarify SZ neurophysiology and guide therapy, single-patient identification remains impractical. This is particularly relevant for treatment-resistant SZ (TRS) whose pathophysiology remains unclear.

**Methods:** Using virtual brain modelling, we simulated large-scale brain dynamics from functional and structural MRI data in 43 healthy controls and 33 SZ patients, stratified into TRS and non-TRS. We used the Wong-Wang model to derive four network parameters reflecting GABAergic and NMDA receptor-mediated glutamatergic transmission: global coupling (G), local inhibitory (J_i_) and excitatory (J_NMDA_) coupling, and recurrent excitation (w_+_). We performed simulations in the whole brain and in resting-state networks, compared model parameters between groups and tested their association with Positive and Negative Syndrome Scale (PANSS) scores.

**Results:** Compared to non-TRS, TRS patients showed increased J_NMDA_ and decreased w_+_ at the whole-brain level, increased J_NMDA_ and G in the dorsal attention network, lower G in the limbic network and increased w_+_ in the visual network. Moreover, J_i_ in the default mode network predicted negative symptom severity while J_NMDA_ in the limbic and ventral attention networks predicted total PANSS scores and disorganisation.

**Conclusions:** These findings suggest that TRS is characterized by network-specific alterations in excitation/inhibition profiles and support the use of personalised multiscale modelling to identify symptom biomarkers.

## Introduction

Schizophrenia (SZ) is a polygenic and multifactorial disorder characterised by a broad spectrum of behavioural, cognitive, affective, sensory and motor symptoms (1), with substantial personal and societal burden (2), posing significant challenges for patients (3) and their families (4): disability-adjusted life years attributable to SZ rose by 65% from 1990 to 2019 (5) and its lifetime suicide rate is estimated at 10% (6).

The diverse clinical manifestations of SZ are increasingly conceptualised across partially dissociable symptom domains, including positive (e.g., delusions and hallucinations), negative (e.g., social withdrawal), disorganisation (e.g., formal thought disorders) and cognitive (e.g., processing speed and working memory deficits) (7). Positive symptoms have traditionally been ascribed to hyperdopaminergia within the mesolimbic pathway and negative and cognitive symptoms to hypodopaminergia within the mesocortical pathway (8). However, contemporary accounts suggest a more distributed pathophysiology, where cognitive and negative symptoms are associated with cortical and cortico-striatal dysfunction resulting from altered glutamatergic and GABAergic signalling (9,10).

Accumulating evidence (1,9,11) points to a major role of the NMDA receptor (NMDA-R) hypofunction (12–14) and distributed anomalies in the excitation/inhibition (E/I) balance, the dynamic interplay between glutamatergic excitation and GABAergic inhibition, in SZ (9,15,16, 17). This is supported by the effects of NMDA antagonists such as ketamine and phencyclidine, which reliably induce psychotomimetic effects in healthy volunteers and exacerbate psychosis in patients (15,16).

Converging electrophysiological, magnetic resonance spectroscopy (MRS) (18), and postmortem (19) evidence further support NMDA-R hypofunction in SZ. Reduced NMDA-R-mediated recruitment of inhibitory interneurons may contribute to cortical disinhibition, altered gamma oscillations and reduced mismatch negativity (10,20,21). Crucially, recent findings suggest a rather specific relationship between cortical E/I imbalance and cognitive deficits in SZ (9,10,22–24), consistent with the limited efficacy of conventional antipsychotics, which act mainly on dopamine D2 receptors and reduce positive but not negative or cognitive symptoms (25,26). Alternative pharmacological strategies have shown early promise, including NMDA-R-enhancing compounds (27,28,29) and muscarinic receptor modulation (30), which allegedly ameliorate negative (31) and cognitive (32) symptomatology.

Resting-state functional magnetic resonance imaging (rs-fMRI) studies indicate that SZ is characterised by large-scale network dysconnectivity, with alterations reported within and between several canonical RSNs (33) comprising the default mode (DMN), frontoparietal (FPN), dorsal attention (DAN), ventral attention (VAN), limbic (LN), somatomotor (SMN) and the visual networks (VN) (34–37). These networks support functions commonly disrupted in SZ: the DMN is implicated in self-referential cognition, the FPN in executive control, the DAN in goal-directed attentional allocation, the VAN in salience detection and attentional reorienting, the LN in affective and mnemonic processing, and the VN and SMN in perceptual integration. Although alterations within these systems have been linked to symptom dimensions (38) and treatment-response variability, findings remain heterogeneous. For example, both hyperconnectivity and hypoconnectivity of the DMN have been reported and associated with different clinical outcomes (34,36,37,39). In contrast, alterations in the DAN appear relatively consistent, with reports of hyperactivity and distinct connectivity profiles differentiating SZ from other psychiatric conditions and predicting cognitive deficits (40–43).

Variability in treatment response further complicates the SZ pathophysiological landscape. Treatment-resistant SZ (TRS) is defined as failure to respond to at least two adequate antipsychotic trials, after which clozapine is the treatment of choice (44). TRS affects approximately one-third of patients and is associated with a more severe clinical profile, including higher PANSS scores, greater cognitive impairment, and higher medication doses (45,46). An imbalance of dopamine-glutamate interplay at postsynaptic density, disrupting cortico-subcortical signalling and synaptic plasticity, has been proposed for TRS (47,48). The superior efficacy of clozapine may relate to its distinct pharmacological profile, characterised by relatively lower D2 affinity and stronger potentiation of NMDA-R-mediated signalling (49), although the use of clozapine entails potentially severe adverse effects such as agranulocytosis and myocarditis (50–52). Consistent with this, TMS (17), multi-omics and post-mortem studies (45) suggest that TRS may represent a distinct neurobiological endophenotype, with alterations extending beyond dopaminergic dysfunction. However, whether these E/I abnormalities are uniformly distributed across the brain or localised to specific RSNs remains unclear.

Although structural and functional alterations have been consistently reported in SZ, conventional connectivity metrics remain descriptive, offering limited insight into the synaptic-level mechanisms underlying network dysfunction and clinical variability. As such, standard approaches often struggle to provide mechanistically interpretable explanations of symptom expression (53,54).

Biophysically grounded virtual brain modelling offers an alternative and promising strategy. By integrating empirical neuroimaging data within generative computational frameworks, platforms such as The Virtual Brain (TVB) enable subject-specific simulations of large-scale neural dynamics with parameters that map onto interpretable neurophysiological processes (55–57). The Wong-Wang neural mass model is especially suited to this aim because it represents each brain region as interacting excitatory and inhibitory neuronal populations, enabling the estimation of biophysically interpretable parameters: global coupling (G), indexing long-range coupling between regions; local inhibitory coupling (J_i_), related to GABAergic control of excitatory neurons; local excitatory coupling (J_NMDA_), encoding NMDA-R-mediated glutamatergic activation of inhibitory interneurons; and recurrent excitation (w_+_), reflecting local excitatory recurrence. This is particularly critical as GABAergic and glutamatergic transmission remain technically challenging to quantify directly and reliably in vivo via MRS, and pharmacological designs probing GABAergic and glutamatergic tone are difficult to scale to large clinical cohorts or resolve at the network level (58).

In this study, we used virtual brain modelling to estimate subject-specific E/I profiles from MRI data at both whole-brain and network levels. We employed the Wong-Wang model to test hypotheses centred on glutamatergic and GABAergic dysfunction. The study aimed to: 1) determine whether E/I profiles distinguished HC, TRS and NON-TRS groups and 2) test whether model parameters predicted positive, negative, disorganisation and global symptom severity.

## Methods and Materials

### Participants

Thirty-three SZ patients (mean age 38.3 ± 9.5 years; age range: 21.2-58.4; 22 men, 11 women) and 43 healthy controls (HC; mean age 39.9 ± 13.4 years; age range: 19.5-75.98; 22 men, 21 women) were recruited at the University of Naples “Federico II”, Outpatient Clinic for Treatment-Resistant Psychoses, Department of Neuroscience, Reproductive Sciences and Odontostomatology.

Patients were stratified into NON-TRS (n=17) and TRS (n=16) subgroups. TRS was defined as failure to respond to at least two adequate antipsychotic trials, whereas NON-TRS patients were classified as responders to non-clozapine antipsychotic treatment. All patients had a previous history of antipsychotic treatment and were receiving antipsychotic medication at the time of assessment. Inclusion criteria for patients were: diagnosis of SZ according to the Diagnostic and Statistical Manual of Mental Disorders (59), established by trained psychiatrists through clinical assessment and review of medical records; age between 18 and 60 years; illness duration longer than two years; no changes in medication or drug dose in the last 6 months; no evidence of worsening of psychotic symptoms in the last three months; absence of brain structural anomalies or other psychiatric or neurological disorders. Clinical stability was assessed by the treating psychiatrist through routine clinical follow-up, clinical interview, medical-record review, and PANSS assessments performed every 3-6 months; the PANSS assessment used in the present study was conducted shortly before MRI acquisition. Exclusion criteria for HC were: history of neurological, psychiatric, or systemic conditions, substance or alcohol dependency, and intake of psychotropic medication during their life. Additionally, HC were screened for history of psychiatric conditions in their family and excluded if they had first-degree relatives with mental disorders (43).

Age at onset, illness duration, medication, chlorpromazine-equivalent dose, substance misuse, handedness and PANSS scores were collected within one month of MRI acquisition. Chlorpromazine-equivalent (60) dose was calculated from the antipsychotic regimen prescribed at the time of clinical/MRI assessment. Age at onset and illness duration are reported by treatment-response subgroup in Table 1. The sample can be considered representative of an established, chronic/relapsing stage of SZ, with medication unchanged for at least six months and chlorpromazine-equivalent doses reflecting treatment at imaging rather than lifetime cumulative exposure.

**Table 1:** Demographic, anamnestic and clinical characteristics of treatment-resistant and non-treatment-resistant patients.

| Variable of Interest | NON-TRS | TRS | Test (df) | *p* |
| --- | --- | --- | --- | --- |
|  | M (SD) | M (SD) |  |  |
| **Age (years)** | 39.2 (9.8) | 37.4 (9.4) | *Wilcoxon W* = 160 | 0.397 |
| **Age of onset (years)** | 24.0 (7.2) | 17.8 (4.4) | *Wilcoxon W* = 216.5 | **0.004\*** |
| **Duration of illness (years)** | 13.65 (7.13) | 17.69 (8.4) | *t* (31) = -1.4922 | 0.146 |
| **Sex (F/M)** | 8/9 (0.51) | 3/13 (0.4) | $\chi^2$ (1) = 1.84 | 0.176 |
| **PANSS<sub>tot</sub>** | 75.44 (13.22) | 92.81 (17.49) | *t* (30) = -3.1699 | **0.003\*** |
| **PANSS<sub>positive</sub>** | 20.53 (5.80) | 24.38 (8.11) | *t* (29) = -1.5068 | 0.143 |
| **PANSS<sub>negative</sub>** | 18.6(6.09) | 26.06 (7.41) | *t* (29) = -3.0517 | **0.005\*** |
| **PANSS<sub>disorganisation</sub>** | 24.81 (7.25) | 31.69 (7.41) | *t* (30) = -2.6528 | **0.013\*** |
| **Chlorpromazine equivalent** | 436.91<br>(282.75) | 534.19<br>(234.13) | *t* (31) = -1.0726 | 0.292 |

The study was conducted in accordance with the ethical principles of the Declaration of Helsinki and approved by the local ethics committee of the University of Naples Federico II and AOU Federico II (protocol No. 195/19). Written informed consent was obtained from all subjects.

### Data acquisition

MRI data were acquired using a Magnetom Trio 3 T scanner (Siemens Healthineers, Erlangen, Germany), equipped with an 8-channel head coil. A T1-weighted volume was acquired using a 3D magnetization prepared rapid acquisition gradient echo (MPRAGE) sequence with the following imaging parameters: TR = 1900 ms; TE = 3.4 ms; TI = 900 ms; flip angle 9°; voxel size 1 x 1 x 1 mm^3^; 160 axial slices. This was used as anatomical reference. Additionally, diffusion-weighted images (DWI) were acquired using a spin echo planar imaging (EPI) sequence with the following parameters: TR = 7400 ms; TE = 88 ms; voxel size = 2.2 x 2.2 x 2.2 mm^3^; 60 slices, with 64 directions at b = 1000 s/mm^2^ in addition to 9 b = 0 s/mm^2^. These images were used for the structural connectivity (SC) analysis. Finally, a T2*-weighted resting-state dataset was acquired using a gradient echo EPI sequence recorded with the following parameters: TR = 2500 ms; TE = 50 ms; voxel size = 3 x 3 x 4 mm^3^; gap = 1 mm; 200 time points; 30 axial slices. This was used for the functional connectivity (FC) analysis.

### MRI preprocessing

MRI data were preprocessed in MATLAB (The MathWorks Inc.) through in-house scripts combining functions from FSL (61), SPM12 (Wellcome Trust Centre for Neuroimaging, Institute of Neurology, University College London), MRtrix3 (62), and BrainSuite (63).

We used rs-fMRI data already preprocessed with slice-time correction, spatial realignment, co-registration to the anatomical image, and normalization to MNI space (43). A CSF mask was created and restricted to regions within the Automatic Lateral Ventricle delIneatioN (ALVIN) mask (64). This was used to remove nuisance regressors (e.g., polynomial trends, head motion) from the preprocessed rs-fMRI data using the component-based noise correction method (65). Finally, the functional images were band-pass filtered (0.008-0.09 Hz).

DWI data were first denoised and corrected for Gibbs ringing artifacts with MRtrix3. Susceptibility-induced distortions were corrected using the T1 anatomical reference image with BrainSuite (63). Finally, eddy current-induced distortions and head motion were corrected with FSL (61).

### Connectome creation

To construct the SC matrix, the brain was parcellated into 124 regions using a registration-based procedure with an ad hoc atlas (see 56) including 78 cortical parcels from the Automated Anatomical Labeling atlas (66), 15 subcortical parcels from FSL-FIRST (67), and 31 cerebellar parcels from the SUIT cerebellar atlas (68). These parcels were used as nodes for subsequent virtual brain modelling.

Structural connectomes were reconstructed from DWI using anatomically constrained tractography, implemented with 30 million streamlines using probabilistic tractography (69). Ipsilateral cerebro-cerebellar connections were removed from the tractograms (see 70). Subject-specific SC matrices, one with weights (i.e., connection strength reflecting the normalized number of streamlines) and one with tract lengths (capturing the average length of connections between regions to account for conduction delays during simulations), were derived by combining the tractogram (71) with the atlas.

Regional BOLD time series were extracted from preprocessed rs-fMRI data by averaging voxel-wise signals within each node. Subject-specific static FC matrices were computed as the Pearson correlation coefficient between pairs of regional time series and subsequently Fisher z-transformed. To characterise the temporal dynamics of FC, dynamic FC (FCD) matrices were also computed (72) using a sliding window of 40 seconds, shifted by one TR across the full time series. For each matrix, we extracted the upper triangular elements and computed pairwise correlations to obtain a time-by-time FCD matrix capturing the evolution of connectivity patterns over time.

To define large-scale functional networks, the nodes of the anatomical atlas were mapped onto canonical resting-state network parcellations derived from previous works (33,73). Based on this mapping, nodes were grouped into seven networks: DMN, FPN, LN, DAN, VAN, VN and SMN. For each subject and for each network, SC, FC and FCD matrices were extracted from the corresponding whole-brain (WB) matrices.

### Virtual brain modelling

We used the TVB platform (74) to simulate large-scale brain dynamics for each participant. Subject-specific SC matrices were used as input, and neural activity in each node was simulated using the Wong-Wang neural mass model (55). The model includes four key parameters: global coupling (G), encoding long-range coupling between regions, local inhibitory coupling (J_i_), reflecting GABAergic synaptic activity, local excitatory coupling (J_NMDA_), capturing glutamatergic synaptic coupling, and recurrent excitation (w_+_), representing recurrent excitation within local excitatory populations. For each node, resting-state BOLD time series were simulated using a Balloon/Windkessel haemodynamic model. Static and dynamic FC matrices were derived.

The four model parameters were optimised in an iterative procedure (56) to maximise the similarity between simulated and empirical FC and FCD. Model goodness of fit was quantified by combining the Pearson correlation coefficient (PCC) between empirical and simulated FC (higher values indicating better fit) and the Kolmogorov-Smirnov (KS) distance, which compares FCD distributions (lower values indicating better fit), into a single cost function equal to ((1-PCC)+KS), where lower values reflect better overall fit.

### Statistical analyses

All statistical analyses were performed using RStudio (Version 2026.01.1; Posit Team, 2026). Demographic, anamnestic, and clinical characteristics of TRS versus NON-TRS patients were compared using t-tests (75), Wilcoxon rank-sum tests (76), or chi-square tests (77) as appropriate according to variable type and assumption assessment. Group differences in model parameters between HCs, TRS and NON-TRS patients were computed at both the WB and network levels using one-way ANOVAs (78) or Kruskal-Wallis tests (79), as appropriate based on assessments of normality and homoscedasticity. When significant omnibus effects were detected, Holm-corrected post-hoc pairwise comparisons were performed to identify specific group differences.

To identify predictors of clinical severity, we performed penalised linear regression using the least absolute shrinkage and selection operator (LASSO) (80). All model parameters were included as candidate predictors, together with relevant covariates (e.g., age at onset and schizophrenia subgroup). LASSO regularisation was used to address potential multicollinearity among predictors and to perform data-driven variable selection. The optimal regularisation parameter (λ) was determined using 10-fold cross-validation. Variables retained by the LASSO model were subsequently entered into standard linear regression models to obtain coefficient estimates and effect sizes. To account for multiple comparisons across regression coefficients, FDR correction (81) was applied. Robustness of results was assessed through sensitivity analysis (see Supplemental Information).

## Results

### Sample characteristics and between-group differences

HCs and SZ patients were well matched for age and biological sex. Sex distribution was comparable across groups (HC: 21F/22M; SZ: 11F/22M). No significant group differences were observed for age (Welch’s t (73.58) = 0.60, p = .553) or biological sex (χ² (1) = 1.26, p = .262).

The SZ subgroups differed only in age of onset, which was entered as a covariate in subsequent linear regression models, and in PANSS total score and negative and disorganisation subscales scores (see Table 1).

### Between-groups differences in E/I profiles

Group comparisons of model-derived parameters across HC, TRS and NON-TRS participants revealed several significant effects (see Figure 2 and Table 2) which were followed by post-hoc pairwise comparisons (see Table S1 for a complete summary of the results).

**Figure 1.**
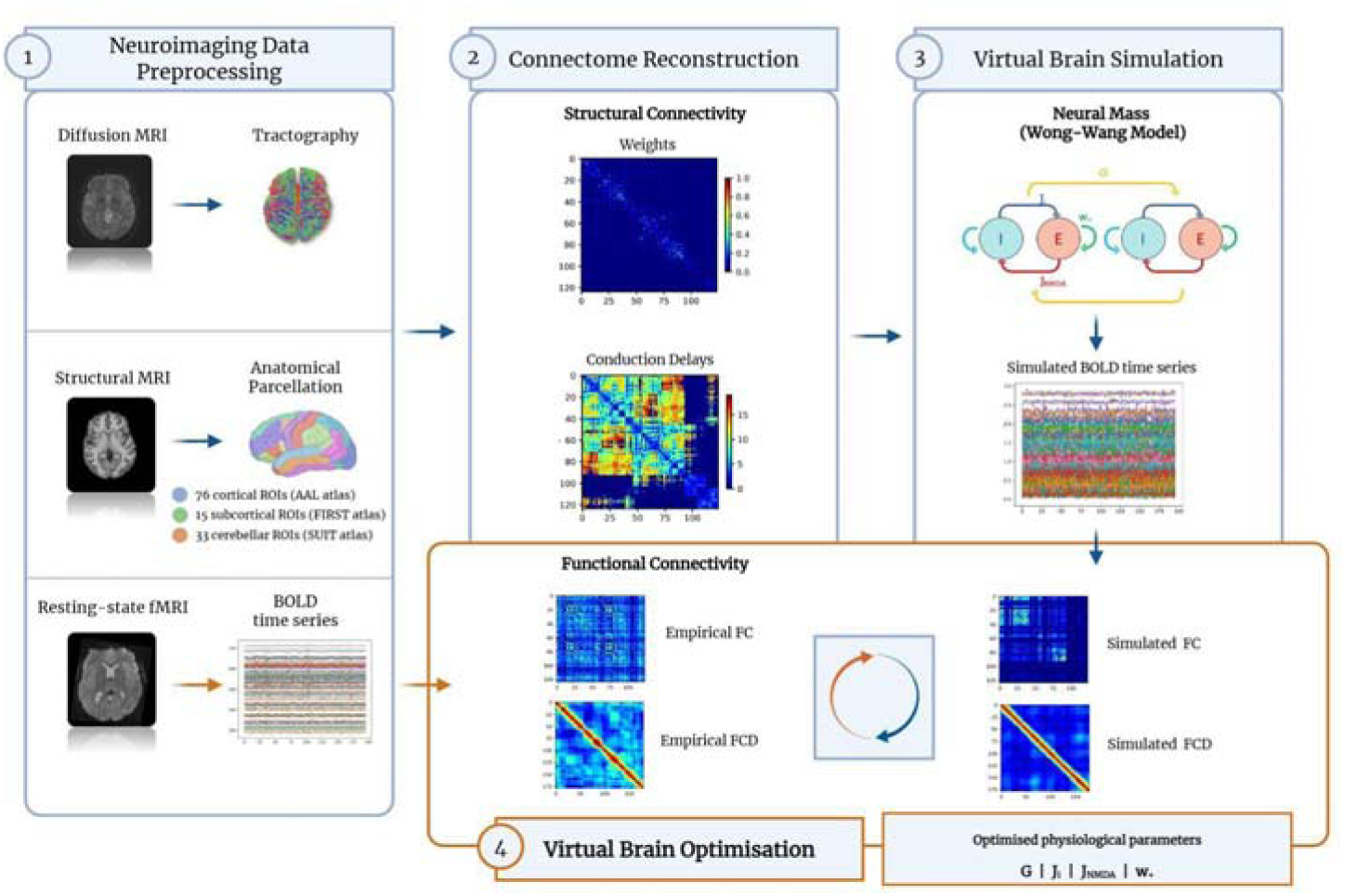
Virtual brain modelling workflow. Structural MRI, diffusion MRI and resting-state fMRI data were preprocessed and used to reconstruct subject-specific brain models. Structural MRI data were used for anatomical parcellation, diffusion MRI for tractography-based structural connectivity reconstruction, and resting-state fMRI for extraction of regional BOLD time series. The brain was parcellated into cortical, subcortical and cerebellar regions, and subject-specific structural connectivity matrices were derived. These matrices were then used as input for The Virtual Brain simulations based on the Wong-Wang neural mass model, which represents each brain region as interacting excitatory and inhibitory neuronal populations. Simulated BOLD time series were generated and used to compute simulated functional connectivity (FC) and dynamic functional connectivity (FCD), which were compared with the corresponding empirical FC and FCD matrices derived from resting-state fMRI. Model parameters were iteratively optimised to maximise the similarity between empirical and simulated FC/FCD, yielding subject-specific estimates of global coupling (G), local inhibitory coupling (Ji), local NMDA-mediated excitatory coupling (JNMDA), and recurrent excitation (w+).

**Figure 2.**
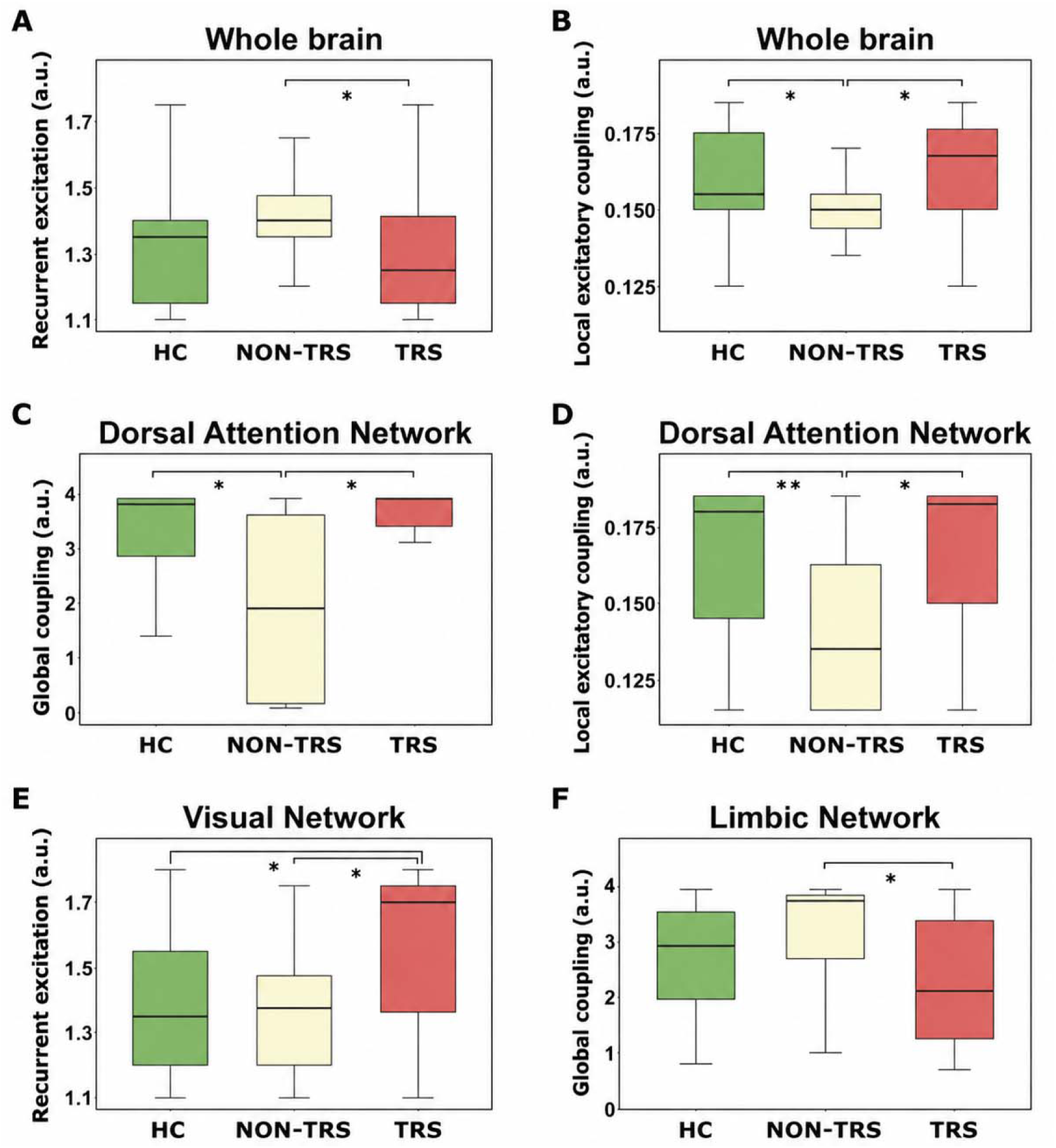
Network-specific group differences in E/I profiles. Boxplots display the median, interquartile range (IQR), and whiskers extending to 1.5 × IQR for whole-brain and resting-state network model parameters derived from the simulations. Model parameters are expressed in arbitrary units (a.u.). Significant differences are indicated with asterisks. At the WB level, w+ (panel A) and JNMDA (panel B) differ between groups. Within the DAN, NON-TRS patients demonstrated reduced G (panel C) and JNMDA (panel D) as compared to both HC and TRS patients. Within the VN, TRS patients exhibited significantly elevated w+ relative to both HC and NON-TRS patients (panel E). Within the LN, NON-TRS patients showed higher G compared to TRS patients (panel F). Together, these findings reveal dissociable, network-specific E/I profiles distinguishing TRS and NON-TRS subgroups. HC = healthy controls; NON-TRS = treatment-responsive schizophrenia; TRS = treatment-resistant schizophrenia; * = p < .05; ** = p < .01.

**Table 2:** Significant between-group differences in model-derived parameters.

| Level /<br>Network | Parameter | HC | NON-TRS | TRS | Omnibus | Omnibus | Significant Holm-corrected post-hoc<br>contrasts |
| --- | --- | --- | --- | --- | --- | --- | --- |
|  |  | M (SD) | M (SD) | M (SD) | test | p |  |
| **Whole brain** | $J_{\text{NMDA}}$ | 0.160 | 0.149 | 0.163 | Kruskal- | .029 | HC vs NON-TRS,<br>pHolm = .044; NON-TRS vs TRS, pHolm =<br>.043 |
|  |  | (0.017) | (0.015) | (0.020) | Wallis |  |  |
| | | | | | $\chi^2(2) =$<br>7.09 | | |
| **Whole brain** | $w_+$ | 1.323 | 1.432 | 1.313 | Kruskal- | .038 | NON-TRS vs TRS,<br>pHolm = .048 |
|  |  | (0.173) | (0.147) | (0.216) | Wallis |  |  |
| | | | | | $\chi^2(2) =$<br>6.54 | | |
| **DAN** | G | 3.009 | 1.900 | 3.131 | Kruskal- | .020 | HC vs NON-TRS,<br>pHolm = .037; NON-TRS vs TRS, pHolm =<br>.025 |
|  |  | (1.374) | (1.583) | (1.399) | Wallis |  |  |
| | | | | | $\chi^2(2) =$<br>7.79 | | |
| **DAN** | $J_{\text{NMDA}}$ | 0.166 | 0.143 | 0.165 | Kruskal- | .005 | HC vs NON-TRS,<br>pHolm = .006; NON-TRS vs TRS, pHolm =<br>.017 |
|  |  | (0.025) | (0.026) | (0.027) | Wallis |  |  |
| | | | | | $\chi^2(2) =$<br>10.60 | | |
| **VN** | $w_+$ | 1.390 | 1.356 | 1.556 | Kruskal- | .022 | HC vs TRS, pHolm =<br>.028; NON-TRS vs<br>TRS, pHolm = .036 |
|  |  | (0.212) | (0.210) | (0.237) | Wallis |  |  |
| | | | | | $\chi^2(2) =$<br>7.65 | | |
| **LN** | G | 2.691 | 3.071 | 2.250 | Kruskal- | .048 | NON-TRS vs TRS,<br>pHolm = .047 |
|  |  | (0.949) | (1.157) | (1.160) | Wallis |  |  |
| | | | | | $\chi^2(2) =$<br>6.08 | | |

At the WB level, NON-TRS patients showed lower J_NMDA_ than both HC and TRS patients, and higher w_+_ than TRS patients. Within the DAN, NON-TRS patients showed lower G and J_NMDA_ relative to both HC and TRS patients. Within the VN, TRS patients showed higher w_+_ than both HC and NON-TRS patients. Finally, within the LN, NON-TRS patients showed higher G than TRS patients.

### Network predictors of clinical severity

Model parameters from all subnetworks, together with relevant clinical and demographic variables, were entered into LASSO regression models to identify predictors of symptom severity. Significant predictors were retained for PANSS total, negative and disorganisation scores, with final model results summarised in Table 3.

**Table 3:** Summary of results from LASSO regression on clinical scores.

| Outcome | Network | Predictors | Effect size | Model | R<sup>2</sup> |
| --- | --- | --- | --- | --- | --- |
| **PANSS** | LN | $J_{\text{NMDA}}$: $\beta = -0.42$, pFDR = **.012** | $\eta^2 p = .29$ | F (2,28) = 9.72, p = .0006 | R<sup>2</sup> = .41; adjR<sup>2</sup> = .37 |
| **Total** | | SZ subgroup (TRS vs NON-TRS): $\beta = 0.42$, pFDR = .012 | $\eta^2 p = .22$ | | |
| **PANSS** | DMN | $J_i$: $\beta = -0.39$, pFDR = .019 | $\eta^2 p = .20$ | F (2,27) = 8.77, p = .0012 | R<sup>2</sup> = .39; adjR<sup>2</sup> = .35 |
| **Negative** | | SZ subgroup (TRS vs NON-TRS): $\beta = 0.49$, pFDR = .0098 | $\eta^2 p = .29$ | | |
| **PANSS** | LN / VAN | $J_{\text{NMDA}}(\text{LIN})$: $\beta = -0.51$, pFDR = .005 | $\eta^2 p = .36$ | F (3,27) = 10.71, p < .001 | R<sup>2</sup> = .54; adjR<sup>2</sup> = .49 |
| **Disorganisation** | | $J_{\text{NMDA}}(\text{VAN})$: $\beta = -0.41$, pFDR = .011 | $\eta^2 p = .29$ | | |
| | | SZ subgroup (TRS vs NON-TRS): $\beta = 0.33$, pFDR = .020 | $\eta^2 p = .18$ | | |

The total PANSS score was significantly predicted by J_NMDA_ in the LN and by the SZ subgroup. Specifically, lower J_NMDA_ values were associated with higher total PANSS scores, indicating that reduced local excitatory coupling within the LN was linked to greater overall clinical severity (Figure 3; Table 3).

**Figure 3.**
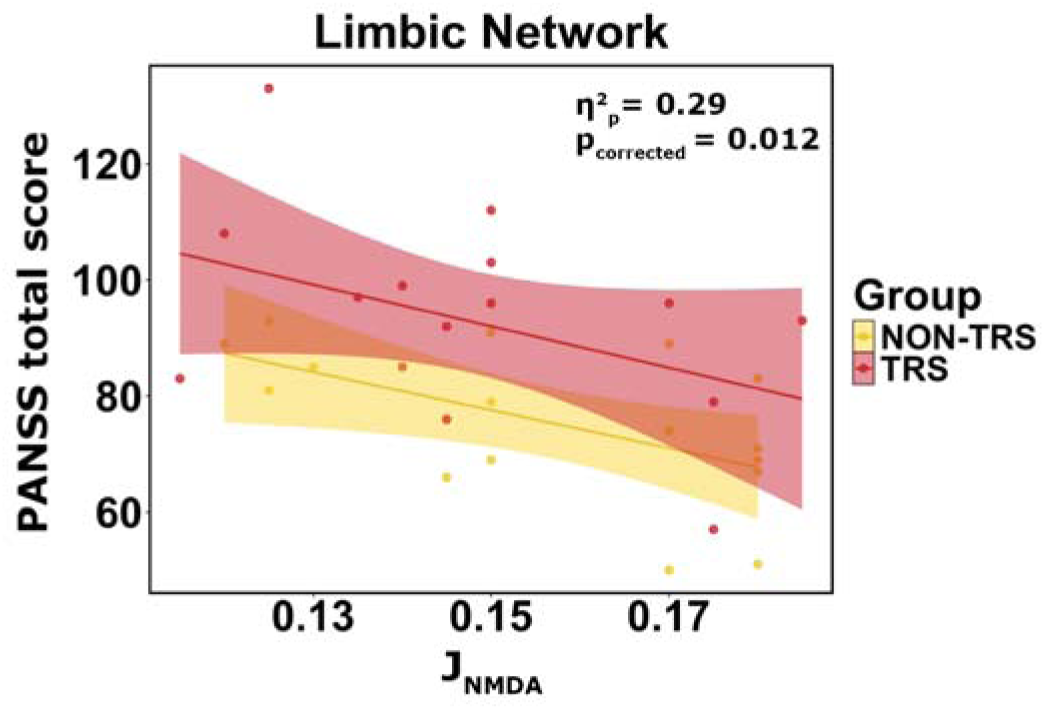
Association between J_NMDA_ in the LN and PANSS total score in SZ patients. Higher values of J_NMDA_ are associated with less severe overall symptomatology. Each point represents a patient, with regression lines shown separately for treatment-resistant schizophrenia (TRS) and NON-TRS groups. The model effect size (partial η²) and the FDR-corrected p-value from the LASSO-selected regression model are displayed.

Negative symptom subscale scores were significantly predicted by J_i_ within the DMN and by the SZ subgroup. Higher local inhibitory coupling values were associated with lower negative symptom severity (Figure 4; Table 3).

**Figure 4.**
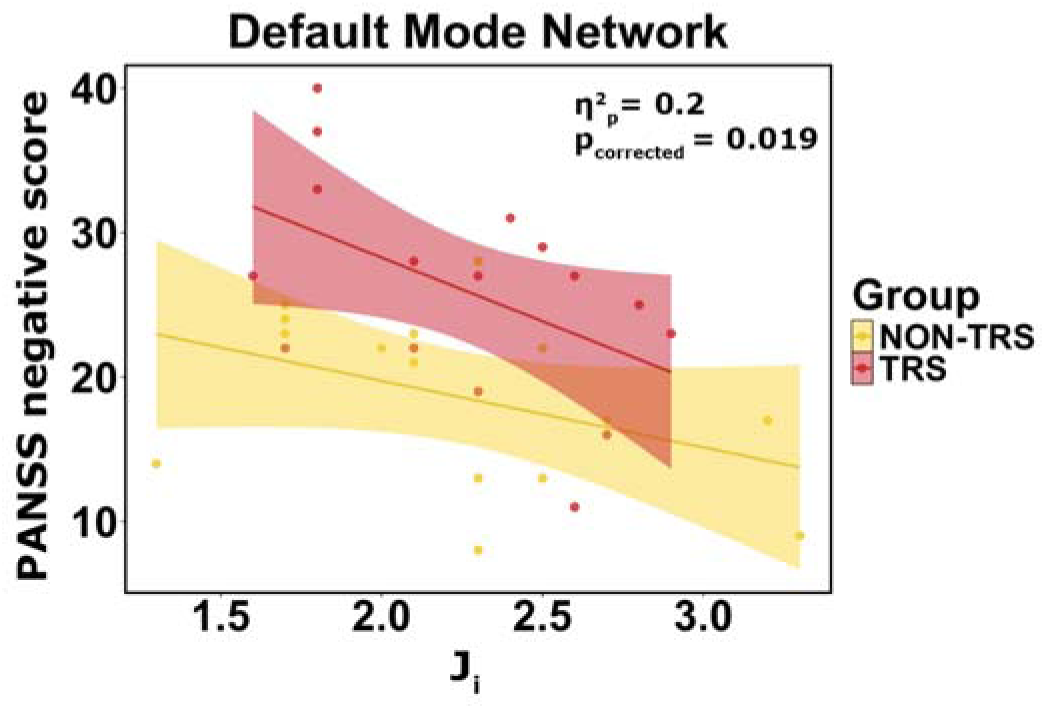
Association between J□ in the DMN and PANSS negative symptom score in SZ patients. Higher values of J_i_ are associated with less severe negative symptoms. Points represent individual observations, with fitted linear trends for TRS and NON-TRS groups. The plot reports the partial η² effect size and FDR-corrected p-value from the regression model.

Symptom severity measured by the disorganisation subscale was significantly predicted by a combination of model parameters in both the LN and the VAN, together with the SZ subgroup (Figure 5; Table 3). Lower J_NMDA_ in the LN (Figure 5, left panel) was associated with greater disorganisation severity. Similarly, reduced J_NMDA_ in the VAN (Figure 5, right panel) predicted higher disorganisation scores.

**Figure 5.**
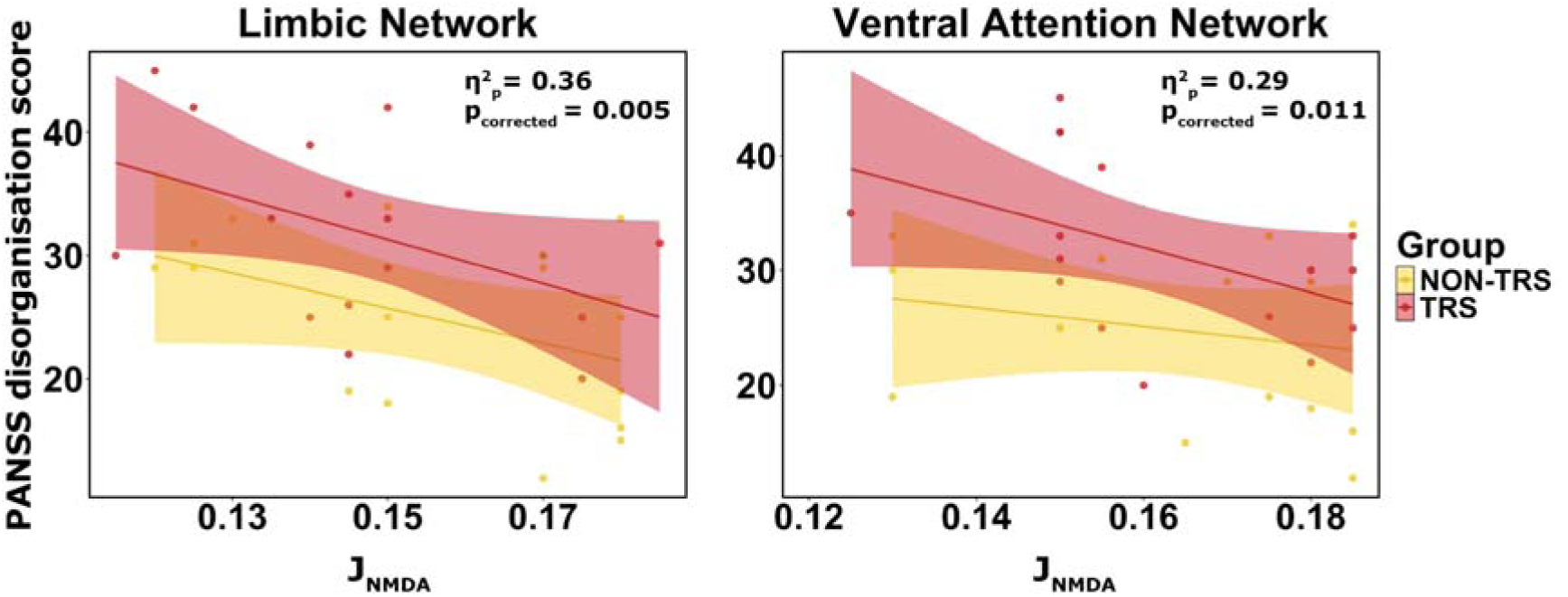
Association between J_NMDA_ in the LN and VAN and PANSS disorganisation scores in SZ patients. Higher values of J_NMDA_ in both LN (left panel) and VAN (right panel) correspond to lower disorganisation symptom severity. Individual observations are shown with group-specific regression lines for TRS and NON-TRS groups. The partial η² effect size and FDR-corrected p-value from the regression analysis are displayed.

## Discussion

Virtual brain modelling revealed distinct, network-specific E/I balance profiles across treatment-response subgroups, with prominent differences between TRS and NON-TRS patients in parameters related to J_NMDA_, w_+_ and G. Group differences emerged at the WB level and within the DAN, VN and LN, while parameters within the DMN, LN and VAN were associated with negative, disorganisation and global symptom severity.

In the DAN, the NON-TRS subgroup showed marked reductions in both G and J_NMDA_ relative to both HC and TRS patients. In the framework of our biophysical model, a lower G corresponds to weaker coupling among nodes within the same network, while lower J_NMDA_ reflects reduced local NMDA-mediated excitatory coupling, consistent with NMDA receptor hypofunction accounts of SZ (12–14). Functionally, these alterations may compromise DAN-mediated attentional control and sensory integration required for goal-directed perception and for resolving ambiguous inputs. The similarity between TRS patients and HC on these metrics highlights subgroup heterogeneity: NON-TRS and TRS may diverge not only in treatment response but also in network-level mechanisms underlying attentional and perceptual dysfunction. Multiple non-mutually exclusive interpretations are plausible: reduced G and J_NMDA_ in the DAN for NON-TRS could reflect an intrinsic, disease-related hypocoupling of attention circuitry contributing to cognitive symptoms (42), modulation by exposure to conventional (non-clozapine) antipsychotics, or compensatory network adaptations linked to clinical improvement.

Additional group differences were observed within other networks. Reduced J_NMDA_ at the WB level in NON-TRS patients compared with both TRS patients and HC may reflect clozapine-induced modulation of NMDA-R-mediated signalling in TRS. However, the cross-sectional design prevents disentangling pharmacological effects from intrinsic differences in excitatory coupling. Conversely, elevated w_+_ in NON-TRS may represent a compensatory upregulation of recurrent excitation in response to reduced NMDA-mediated coupling.

In the VN, TRS patients exhibited higher w_+_ than both HC and NON-TRS patients, suggesting enhanced recurrent excitatory dynamics within VN nodes. This finding is consistent with abnormalities in early-stage visual processing documented in both chronic SZ and antipsychotic-naïve first-episode psychosis, suggesting that VN dysfunction emerges in early stages of the disease (82–84). Recent work further suggests that occipital and fronto-limbic-occipital connectivity patterns may differentiate TRS from NON-TRS patients, supporting the potential relevance of VN alterations for biological stratification (85–88). Notably, a recent PET study showed that brain glucose metabolism in the fusiform gyrus distinguishes TRS patients from both NON-TRS patients and HCs (89). However, the functional significance of this result should be clarified in future studies combining virtual brain modelling with behavioural or task-based measures of sensory processing.

Furthermore, NON-TRS patients exhibited higher LN G than TRS ones, suggesting stronger intrinsic coordination and more coherent integration of regions related to emotions and memory, potentially preserving affective regulation and responsiveness to conventional antipsychotic treatment (90). Conversely, reduced limbic global coupling in TRS points to weaker intrinsic synchronisation within the limbic system, consistent with a less stable or less efficiently coordinated network state.

Our LASSO regression analyses revealed network-specific relationships between estimated E/I dynamics and symptom dimensions, after accounting for treatment-response subgroup. First, J_i_ at the DMN level emerged as a robust predictor of negative symptoms: reduced J_i_, interpreted as weakened GABAergic/inhibitory control, was associated with more severe negative symptomatology. This observation is consistent with the notion that deficient interneuron control permits excessive, internally-driven DMN activity, which, in turn, may manifest clinically as affective flattening, avolition, and social withdrawal (91–93). Specifically, weakened inhibitory control in the DMN could bias information processing toward self-referential, internally generated activity at the expense of outward-directed engagement. Second, reduced J_NMDA_ in the LN was associated with higher total PANSS scores and greater disorganisation severity, suggesting that limbic NMDA-R-dependent signalling may contribute to the broader clinical burden of SZ (94,95). Finally, disorganisation symptoms were also associated with reduced J_NMDA_ within the VAN, suggesting that NMDA-related mechanisms spanning limbic and attentional circuits may contribute to cognitive fragmentation and impaired integration of information across large-scale networks. No significant predictors of positive symptoms were identified. This finding is in line with the general notion that typical antipsychotics, with a strong affinity for D2 receptors, mainly ameliorate positive symptomatology, with limited benefit for negative or cognitive symptoms (25). This null result may also reflect the specific focus of the computational model, which captures glutamatergic and GABAergic dynamics but does not explicitly model dopaminergic signalling, widely considered a key neurochemical driver of positive symptoms. Future extensions incorporating dopamine-sensitive mechanisms may therefore better capture variability in positive symptom expression (54). Across the significant models, subgroup membership also contributed to symptom severity, emphasising the importance of treatment-response status when interpreting associations between model parameters and clinical outcomes.

Taken together, the group differences and symptom associations suggest that treatment-response condition and symptom dimensions are associated with partially distinct network-level mechanisms. Functionally interpretable parameters derived from subject-specific multiscale models, such as J_i_ and J_NMDA_, may help bridge macroscopic connectivity phenotypes with putative mechanisms and generate at the level of synapses and circuits hypotheses relevant to targeted pharmacological or neuromodulatory strategies. These implications nevertheless require validation in independent and longitudinal cohorts.

Some limitations should be considered when interpreting the present findings. First, causal inference is limited by the cross-sectional design and by potential confounds related to medication, illness duration and sample size. Medication exposure, particularly clozapine treatment in TRS, cannot be disentangled from intrinsic disease-related differences. Second, model parameters were inferred from a generative model rather than measured directly in vivo and should therefore be considered indirect proxies of E/I dynamics rather than direct measures of glutamatergic or GABAergic neurotransmission. Third, the present findings were not validated in independent datasets and should therefore be interpreted as exploratory and hypothesis-generating.

Addressing these limitations will require validation in independent cohorts, longitudinal modelling in early-stage or minimally treated patients, and multimodal validation using MRS, PET, TMS or behavioural measures. Experimental perturbations, including in silico lesioning or parameter sweeps, could further test whether the identified parameter changes are sufficient to reproduce the connectivity signatures observed in patients. In the longer term, such approaches may contribute to the development of personalised computational models, or “brain digital twins”, aimed at characterising individual pathophysiology and simulating candidate pharmacological or neuromodulatory interventions. At present, however, this remains a theoretical perspective in clinical psychiatry and should be considered a future research direction rather than a direct clinical implication of the present study.

The broader implication of this framework is that generative modelling may provide a way to move from the description of abnormal functional connectivity to experimentally testable hypotheses about the circuit-level mechanisms underlying clinically relevant phenotypes. One possible mechanistic interpretation is that altered NMDA receptor-dependent signalling may affect the recruitment or modulation of inhibitory interneurons, thereby shifting E/I balance within selectively affected networks. This interpretation remains necessarily tentative, as the present model parameters are indirect proxies rather than direct measures of synaptic neurotransmission. Nevertheless, the ability to relate subject-specific model parameters to symptom dimensions and treatment-response phenotypes provides a basis for investigating whether apparently heterogeneous SZ presentations can be understood in terms of distinct combinations of network-level circuit alterations.

In this context, subject-specific virtual brain models could offer a framework for testing mechanistic hypotheses about patient heterogeneity and, in the long term, for simulating candidate pharmacological or neuromodulatory interventions. Such applications remain prospective and require replication in larger independent cohorts, longitudinal validation, and integration with complementary biological measures.

## Acknowledgments

This work was supported by #NEXTGENERATIONEU (NGEU) and funded by the Ministry of University and Research (MUR); National Recovery and Resilience Plan (NRRP) project; MNESYS (PE0000006) – A Multiscale integrated approach to the study of the nervous system in health and disease (DN. 1553 11.10.2022) and EBRAINS-Italy (PIR0000011, CUP B51E22000150006, “EBRAINS-Italy”) awarded to EDA and CAMGWK, and the EU Horizon Europe program projects EBRAINS2.0 (101147319), Virtual Brain Twin (101137289) and TEF-Health (101100700) awarded to EDA. CAMGWK also receives funding from UKRI (STFC #UKRI4203), BRC (#BRC704/CAP/CGW), MRC (#MR/S026088/1), Ataxia UK, Rosetrees Trust (#PGL22/100041 and #PGL21/10079).

## Supplemental Information

### Sensitivity analyses

For each linear regression model (see section “Network Predictors of Clinical Severity”), potentially influential observations were detected using studentized residuals, Cook’s distance and leverage values. The models were refitted after excluding flagged observations.

Regression diagnostics did not identify residual outliers based on studentized residuals. Although some observations exceeded Cook’s distance and/or leverage thresholds, sensitivity analyses yielded comparable effects and did not alter the interpretation of the main predictors (see Table S2).

## Table Legends

**Table 1 Values are reported as mean (and SD), except for sex, which is reported as female/male (female proportion).** Between-group comparisons were performed using independent-samples t-tests, Wilcoxon rank-sum tests, or chi-square tests, as indicated in the table. PANSS = Positive and Negative Syndrome Scale; CPZ = chlorpromazine equivalents; M = mean; SD = standard deviation; χ² = chi-square test. *p < .05.

**Table 2 The table summarises model parameters showing significant omnibus group effects followed by significant Holm-corrected post-hoc pairwise comparisons across HC, TRS, and NON-TRS patients.** For each parameter, the table reports group mean (SD), the WB level or resting-state network, the omnibus test statistic and p-value, and the significant post-hoc contrasts. Only effects surviving Holm-corrected post-hoc testing are reported. G = global coupling; JNMDA = local NMDA-mediated excitatory coupling; w+ = recurrent excitation; DAN = dorsal attention network; VN = visual network; LN = limbic network; M = mean; SD = standard deviation.

## Supplemental Information

**Table S1:** Summary of omnibus test statistics for between-group comparisons of TVB-derived parameters.

| Level / Network | G | $J_i$ | $J_{\text{NMDA}}$ | $w_+$ |
| --- | --- | --- | --- | --- |
| **WB** | $F(2,73) = 0.36, p = .698$ | $F(2,73) = 0.17, p = .843$ | $\chi^2(2) = 7.09, p = .029$ | $\chi^2(2) = 6.54, p = .038$ |
| **DMN** | $\chi^2(2) = 0.20, p = .904$ | $F(2,73) = 0.86, p = .427$ | $\chi^2(2) = 0.28, p = .871$ | $\chi^2(2) = 0.29, p = .866$ |
| **FPN** | $F(2,73) = 1.80, p = .173$ | $\chi^2(2) = 1.31, p = .519$ | Welch $F(2,29.17) = 1.06, p = .359$ | Welch $F(2,30.44) = 1.07, p = .357$ |
| **SMN** | $\chi^2(2) = 0.33, p = .849$ | $\chi^2(2) = 0.57, p = .754$ | $\chi^2(2) = 0.53, p = .768$ | $\chi^2(2) = 0.65, p = .723$ |
| **VAN** | $\chi^2(2) = 2.25, p = .325$ | $F(2,73) = 1.32, p = .274$ | $\chi^2(2) = 0.97, p = .616$ | $\chi^2(2) = 0.58, p = .746$ |
| **VN** | $\chi^2(2) = 0.46, p = .796$ | $\chi^2(2) = 0.39, p = .822$ | $\chi^2(2) = 5.70, p = .058$ | $\chi^2(2) = 7.65, p = .022$ |
| **DAN** | $\chi^2(2) = 7.79, p = .020$ | $\chi^2(2) = 4.30, p = .117$ | $\chi^2(2) = 10.60, p = .005$ | $\chi^2(2) = 3.39, p = .184$ |
| **LN** | $\chi^2(2) = 6.08, p = .048$ | $\chi^2(2) = 6.14, p = .046$ | $\chi^2(2) = 0.97, p = .615$ | $\chi^2(2) = 1.47, p = .480$ |

**Table S2:**
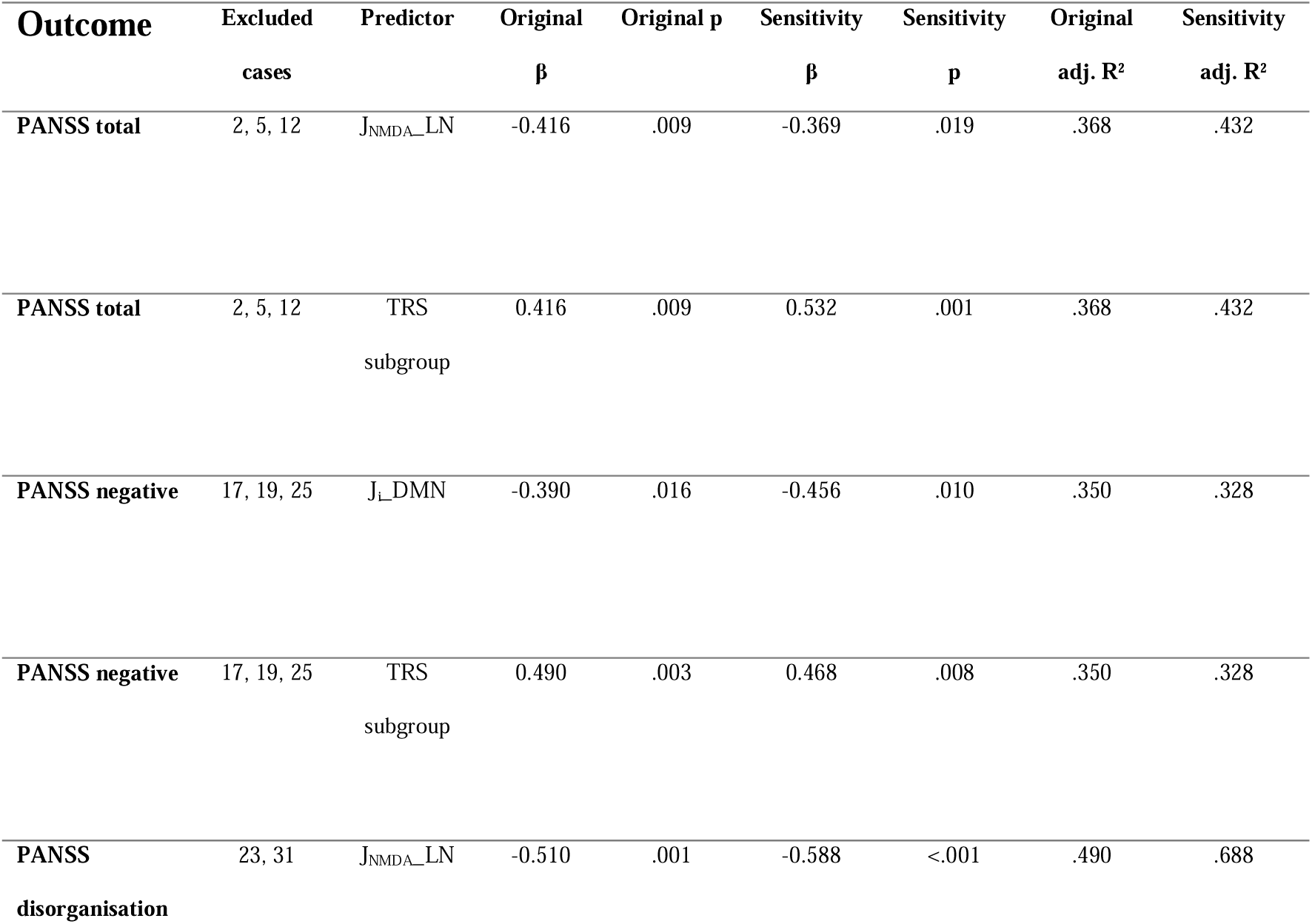
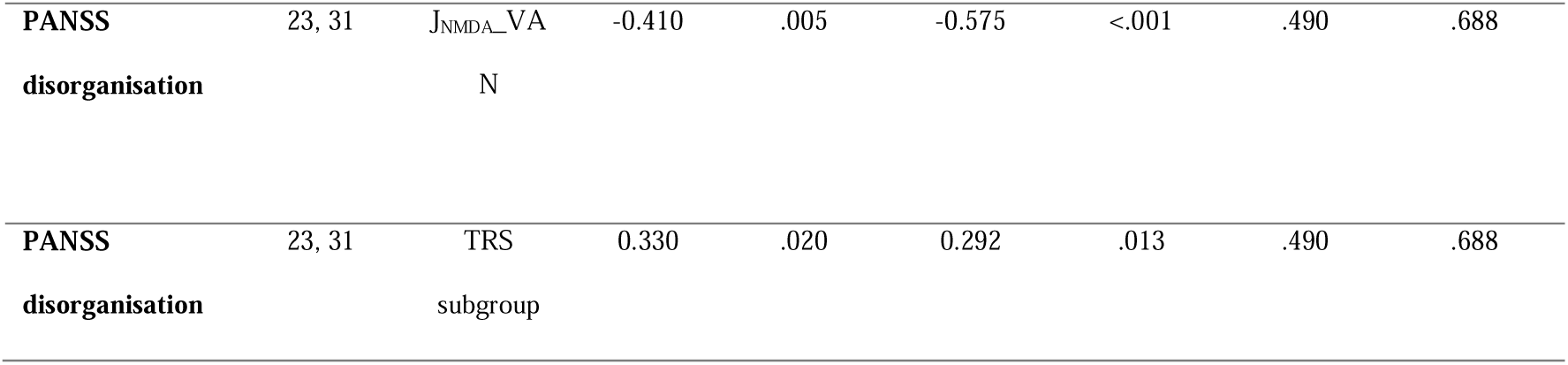
Summary of results from sensitivity analyses on regression models.

